# Highly expressed cell wall genes contribute to robustness of sepal size

**DOI:** 10.1101/2024.10.14.618238

**Authors:** Diego A. Hartasánchez, Mathilde Dumond, Nelly Dubrulle, Françoise Monéger, Arezki Boudaoud

## Abstract

Reproducibility in organ size and shape is a fascinating trait of living organisms. The mechanisms underlying such robustness remain, however, to be elucidated. Taking the sepal of Arabidopsis as a model, we investigated whether variability of gene expression plays a role in variation of organ size and shape. Previous work from our team identified cell-wall related genes as being enriched among the genes whose expression is highly variable. We then hypothesized that the variation of measured morphological parameters in cell-wall related single knockout mutants could be correlated with the variation in gene expression of the corresponding gene (the knocked-out gene) in wild-type plants. We analyzed sepal size and shape from 16 cell-wall mutants and found that sepal size variability correlates positively, not with gene expression variation, but with mean gene expression of the corresponding gene in wild type. These findings support a contribution of cell-wall related genes to the robustness of sepal size.

## Introduction

Organisms of the same species typically exhibit a remarkably reproducible morphological development despite high variability at the cellular level. The invariant expression of phenotype in the face of environmental and/or genetic perturbations, commonly referred to as “robustness” (Félix & Barkoulas, 2015), is indeed an important characteristic of living beings (Schmalhausen, 1949; Waddington, 1953, 1959). In plants, molecular mechanisms have been found to modulate morphogenetic robustness to environmental perturbations, as in the case of heat-shock proteins (Queitsch et al., 2002), and to genetic changes such as whole genome duplications (Lachowiec et al., 2016). Robustness, however, also refers to developmental stability despite systemic internal noise (Alvarez-Buylla et al., 2008). Indeed, gene expression has an important stochastic component attributed to a combination of external and internal noise, as initially shown in bacteria (Elowitz et al. 2002). In a multicellular context, such as that of Arabidopsis plants, gene expression appears to be extremely variable in time and space (Joseph et al., 2015; Araújo et al., 2017; Meyer et al., 2017). In fact, when measuring variability in gene expression at the whole-organism level, there are some genes that exhibit very high variability between individuals. This variability itself has been observed to differ between day and night, for example, in Arabidopsis seedlings (Cortijo et al. 2019) and between developmental stages in *C. elegans* (Zalts & Yanai, 2017) and Drosophila (Liu et al., 2020). Development is, then, not only robust to gene expression variability, but also, possibly dependent on it.

In a recent paper, Hartasánchez et al. (2023) used the sepal of wild-type Arabidopsis plants to identify modules of co-expressed genes which co-vary with sepal morphology. Cell-wall related genes were found to be over-represented in two of these modules. In addition, highly variable genes were also enriched in cell-wall related genes. Building upon these results, we wanted to check if cell-wall related genes could be involved in the robustness of sepal morphology. We selected a sample of 16 genes, and we studied the corresponding mutants to evaluate the variability of their sepal size and shape in relation to the variability of expression of the corresponding gene in the wild-type sepal. Our results reveal a positive correlation between the level of expression of the genes in wild type and variability of size in the corresponding mutants. Altogether, our work supports a contribution of highly expressed cell-wall related genes to the robustness of sepal size.

## Materials and methods

### Plant Material

Col-0 *Arabidopsis thaliana* plants were grown on soil at 20°C in short day conditions (8 h light/16 h dark) for 20 days before being transferred to long day conditions (16 h light/8 h darkness). Sepals were dissected from secondary inflorescences after at least 10 siliques were formed. We assessed the final shape of sepals (stage 13, according to Smyth et al. 1990).

### Mutants

Mutant seeds (from Col-0 background) were obtained from well-characterized stocks (Desnos et al., 1996; Bringmann et al., 2012; Sénéchal, 2013; Endler et al., 2015) or from SAIL and SALK collections maintained at NASC (McElver et al., 2001; Alonso et al., 2003) as described in Supplementary Table 1. Mutant plants were genotyped following O’Malley et al. (2015), except for *csi1-3* and *prc1* which have distinctive phenotypes.

### Extraction of morphological parameters from mutant lines

We analyzed sepal contours and quantified area, length, width, and aspect ratio. We only analysed these four geometrical parameters because they were the only parameters in common with the study relating gene expression to 3D sepal morphology (Hartasánchez et al., 2023). Briefly, we flattened the sepals between two slides and took photographs with a black background under a dissecting microscope, following Hong et al. (2016). We used Python scripts (Van Rossum & Drake, 2009) to segment and align sepals, and to extract morphological parameters (Hong et al., 2016).

### Data analysis

Raw data consisting of measurements of length, width, area, and aspect ratio for control plants (Supplementary Table 2; 11 control batches) and for each mutant (Supplementary Table 3; 16 mutants) were analyzed. The difference between mutant and wild type (from the corresponding control batch) for each morphological parameter was calculated as: [mean(mutantParameter) - mean(controlBatchParameter)] / mean(controlBatchParameter). Since we are interested in the magnitude of this difference and not its sign, we used this difference in absolute numbers when testing for correlations. The squared coefficient of variation (CV^2^) for each parameter was calculated as: [sd(mutantParameter) / mean(mutantParameter)] ^2. The average gene expression for each mutant gene in wild type was obtained from Hartasánchez et al. (2023) (Supplementary Table 4). Source data for Figures 1 & 2 are shown in Supplementary Table 5.

**Figure 1.**
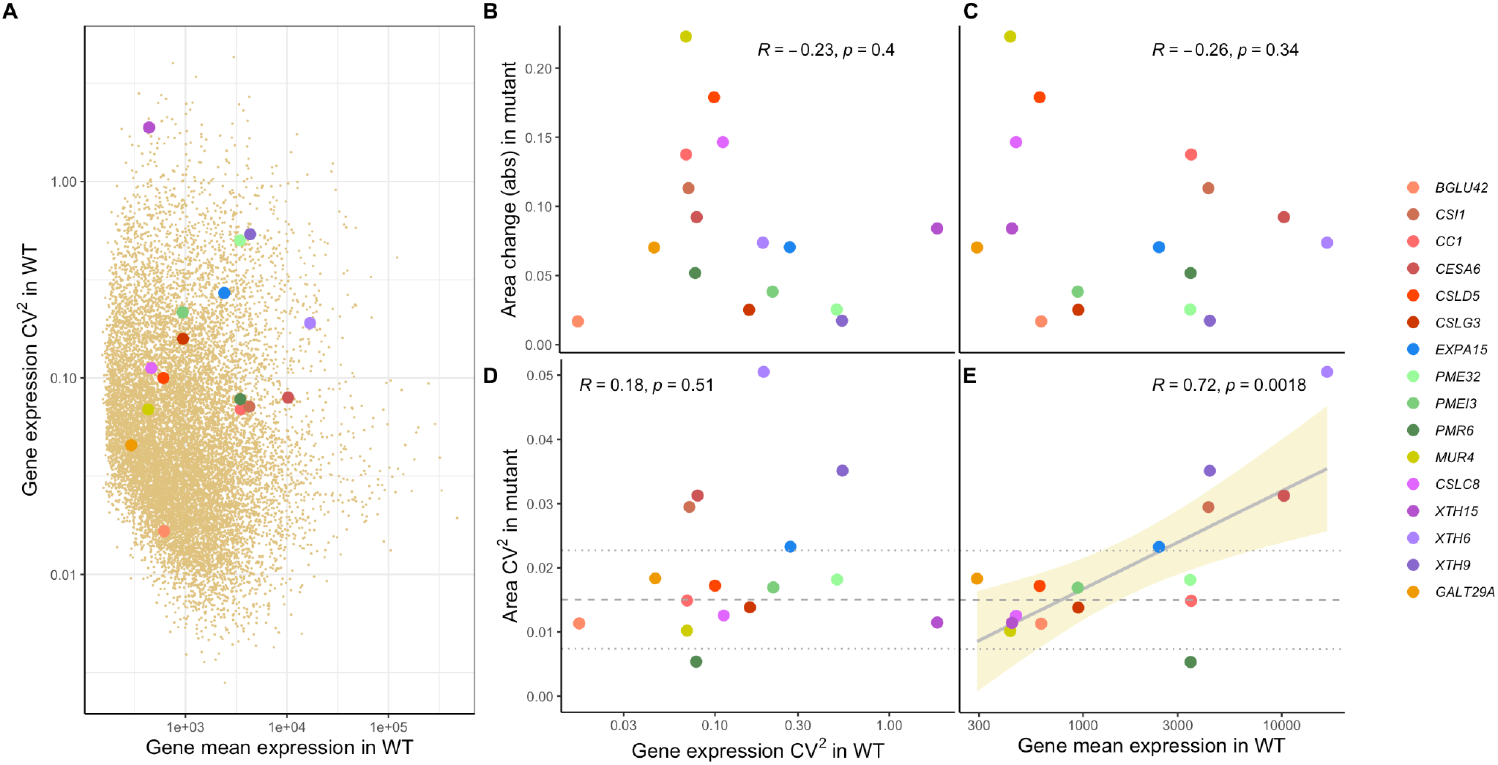
A) Squared coefficient of variation (CV^2^) of gene expression against mean gene expression in wild-type (WT) Col-0 sepals for 16 mutant genes (in colors as shown in the legend) over the corresponding values for 14,085 genes expressed in sepals (from Hartasánchez et al., 2023; dark tan points). B & C) Effect (in absolute value, denoted ‘abs’) of the knockout mutation of each gene on sepal area (corrected by sepal area in wild-type plants for each batch) against gene expression CV^2^ in WT (B), and against gene mean expression in WT (C). D & E) CV^2^ of sepal area in knockout mutants against gene expression CV^2^ in WT (D), and against gene mean expression in WT (E). Each colored point corresponds to one gene. Pearson correlation coefficient R and p-values are shown on the top in each plot. Gray solid line shows linear model adjustment with standard error in tan shade. Gray dashed lines correspond to average CV^2^ in area measurements in WT Col-0 control batches with dotted lines showing average plus/minus one standard deviation.

### Mutant subsampling

We ran 1000 subsampling replicates. In each replicate, we extracted a random sample of 30 sepals (without replacement) for each mutant. For each parameter and each replicate, we tested if there was a correlation between the parameter’s CV^2^ (obtained from the 30 subsamples of each mutant) and the log mean gene expression for the corresponding gene in wild type. We obtained Pearson correlation coefficients (R) and p-values for the 1000 correlation tests for each parameter.

### Leave x-out experiments

We obtained Pearson correlation coefficients (R) and corresponding p-values for Area, Length and Width CV^2^ values against gene mean expression in wild type across a subselection of mutants. Leave 1-out experiments tested 16 correlations, leaving one mutant out at a time. Leave 2-out and leave 3-out experiments tested 240 and 3360 correlations, respectively, accounting for all possible combinations in which two or three mutants were left out. The leave *pmr6*+x experiments consisted in eliminating the *PMR6* mutant from the list and then performing the leave x-out experiments, with 15, 210 and 2730 combinations tested for *pmr6*+1, *pmr6*+2 and *pmr6*+3, respectively.

## Results

To explore the link between cell-wall related gene expression and robustness of sepal morphology, we decided to exploit an unpublished mutant dataset previously produced by our team (Dumond, 2017). This mutant dataset had been generated in the context of sepal phenotype exploration and contained data for 16 of the 718 cell-wall related genes expressed in sepal (Table 1; Supplementary Table 1). The mutants had been selected based on the following features: involvement of the corresponding gene in synthesis or remodelling of cell wall components, relatively higher expression of the corresponding gene in sepals compared to other organs at stage 12 (Schmid et al., 2005), and availability of mutants with T-DNA insertions in exons (we considered one mutant allele for each of these genes). The 16 corresponding genes can be grouped according to their (putative) functions as follows (Table 1): six genes encoding proteins related to cellulose [one hydrolase (BETA GLUCOSIDASE 42), two interactors of the cellulose synthase complex (CELLULOSE SYNTHASE-INTERACTIVE PROTEIN 1 and COMPANION OF CELLULOSE SYNTHASE 1), and three cellulose synthases (CELLULOSE SYNTHASE 6, CELLULOSE SYNTHASE-LIKE D5, CELLULOSE SYNTHASE-LIKE G3)]; one gene encoding a protein related to cellulose and hemi-cellulose [an expansin (EXPA15)]; three genes encoding proteins related to pectin [a pectin methylesterase (PECTIN METHYLESTERASE 32), a pectin methylesterase inhibitor (PECTIN METHYLESTERASE INHIBITOR 2), and a pectate lyase (POWDERY MILDEW RESISTANT 6)]; one gene encoding a protein related to pectin and hemicellulose [a protein involved in synthesis of nucleotide-arabinose and necessary for arabinosylation of pectin and hemicellulose (MURUS 4)]; four genes encoding proteins related to hemicellulose [one glucan synthase (CELLULOSE SYNTHASE-LIKE C8), and three xyloglucan endotransglucosylase / hydrolases (XTH15, XTH6, and XTH9)]; and one gene encoding a protein related to arabinogalactan proteins [a glycosyltransferase (GLYCOSYLTRANSFERASE 29A)].

**Table 1.**

| GENE | GENE ID | RELATED TO | INVOLVED IN or ENABLES |
| --- | --- | --- | --- |
| *BETA GLUCOSIDASE 42*<br>(*BGLU42*) | AT5G36890 | CELLULOSE | cellulose catabolic process |
| *CELLULOSE SYNTHASE-INTERACTIVE PROTEIN 1*<br>(*CSI1*) | AT2G22125 | CELLULOSE | cellulose biosynthetic process |
| *COMPANION OF CELLULOSE SYNTHASE 1*<br>(*CC1*) | AT1G45688 | CELLULOSE | linking cellulose synthase complex and microtubules |
| *CELLULOSE SYNTHASE 6*<br>(*CESA6*) | AT5G64740 | CELLULOSE | cellulose biosynthetic process |
| *CELLULOSE SYNTHASE-LIKE D5*<br>(*CSLD5*) | AT1G02730 | CELLULOSE | cellulose biosynthetic process |
| *CELLULOSE SYNTHASE-LIKE G3*<br>(*CSLG3*) | AT4G23990 | CELLULOSE | cellulose biosynthetic process |
| *EXPANSIN A15*<br>(*EXPA15*) | AT2G03090 | CELLULOSE/<br>HEMICELLULOSE | cell wall organization |
| *PECTIN METHYLESTERASE 32*<br>(*PME32*) | AT3G43270 | PECTIN | pectin catabolic process |
| *PECTIN METHYLESTERASE INHIBITOR 3*<br>(*PMEI3*) | AT5G20740 | PECTIN | negative regulation of pectin catabolic process |
| *POWDERY MILDEW RESISTANT 6*<br>(*PMR6*) | AT3G54920 | PECTIN | pectin catabolic process |
| *MURUS 4*<br>(*MUR4*) | AT1G30620 | PECTIN/<br>HEMICELLULOSE | galactose metabolic process<br>for cell wall biogenesis |
| *CELLULOSE SYNTHASE-LIKE C8*<br>(*CSLC8*) | AT2G24630 | HEMICELLULOSE | cell wall organization |
| *XYLOGLUCAN ENDOTRANSGLUCOSYLASE/HYDROLASE 15*<br>(*XTH15*) | AT4G14130 | HEMICELLULOSE | xyloglucan metabolic process |
| *XYLOGLUCAN ENDOTRANSGLUCOSYLASE/HYDROLASE 6*<br>(*XTH6*) | AT5G65730 | HEMICELLULOSE | xyloglucan metabolic process |
| *XYLOGLUCAN ENDOTRANSGLUCOSYLASE/HYDROLASE 9*<br>(*XTH9*) | AT4G03210 | HEMICELLULOSE | xyloglucan metabolic process |
| *GLYCOSYLTRANSFERASE 29A* (*GALT29A*) | AT1G08280 | ARABINOGALACTAN<br>PROTEINS | protein glycosylation |

Because it is difficult to compare developmental stages between mutant and wild-type plants, we focused on the size and shape of sepals that had ceased growing (stage 13). For each mutant and their corresponding wild-type controls (Supplementary Table 2), we had obtained length, width, area, and aspect ratio with samples ranging from 39 to 90 sepals (see Materials and Methods & Supplementary Table 3). For the level of expression of the corresponding genes in wild-type sepals, we reasoned that transcriptome of sepals at stage 11 would be a good predictor of the final size of sepals (stage 13) because stage 11 precedes growth cessation. Accordingly, we used the data generated by Hartasánchez et al. (2023) composed of transcriptomes of 27 individual sepals (stage 11) from Col-0 background generated by bulk RNA-seq (Supplementary Table 4). Despite cell-wall related genes being enriched in highly variable genes (Hartasánchez et al., 2023), the genes represented within our set of mutants are widespread in both gene expression CV^2^ and mean gene expression levels, as shown in Figure 1A, in comparison with the 14,085 genes expressed in sepals from Hartasánchez et al. (2023).

We first hypothesized that the level of expression of a given gene in wild-type plants could predict the effect on morphology of the corresponding knockout mutant. We thus examined if knockout mutants of genes with higher expression in wild-type plants exhibited a stronger phenotype (difference in morphological parameters compared to wild type; see Materials and methods) than genes with lower expression. Our data (Supplementary Table 5), however, shows no correlation between the size or shape of mutant lines and the level of expression of the corresponding gene in wild-type plants (Figure 1C).

We then hypothesized that genes with higher expression variability in wild type would tend to be more important for robust morphogenesis, and hence, when knocked out, would have a stronger effect on sepal size and shape or their variability. Again, this was not supported by the data (Figure 1B & 1D; Supplementary Table 5). Finally, we hypothesized that knocking down highly expressed genes in wild type would prove more difficult to cope with than knocking down genes with lower expression. Accordingly, mutant phenotypic variability would correlate with the level of gene expression in wild type. Indeed, there is a significant correlation between the coefficient of variability (CV^2^) of the mutant phenotypes (for area, length and width, but not aspect ratio) and the log mean expression in the wild type of the corresponding knocked out genes (Figure 1E & Figure 2), supporting the latter hypothesis.

**Figure 2.**
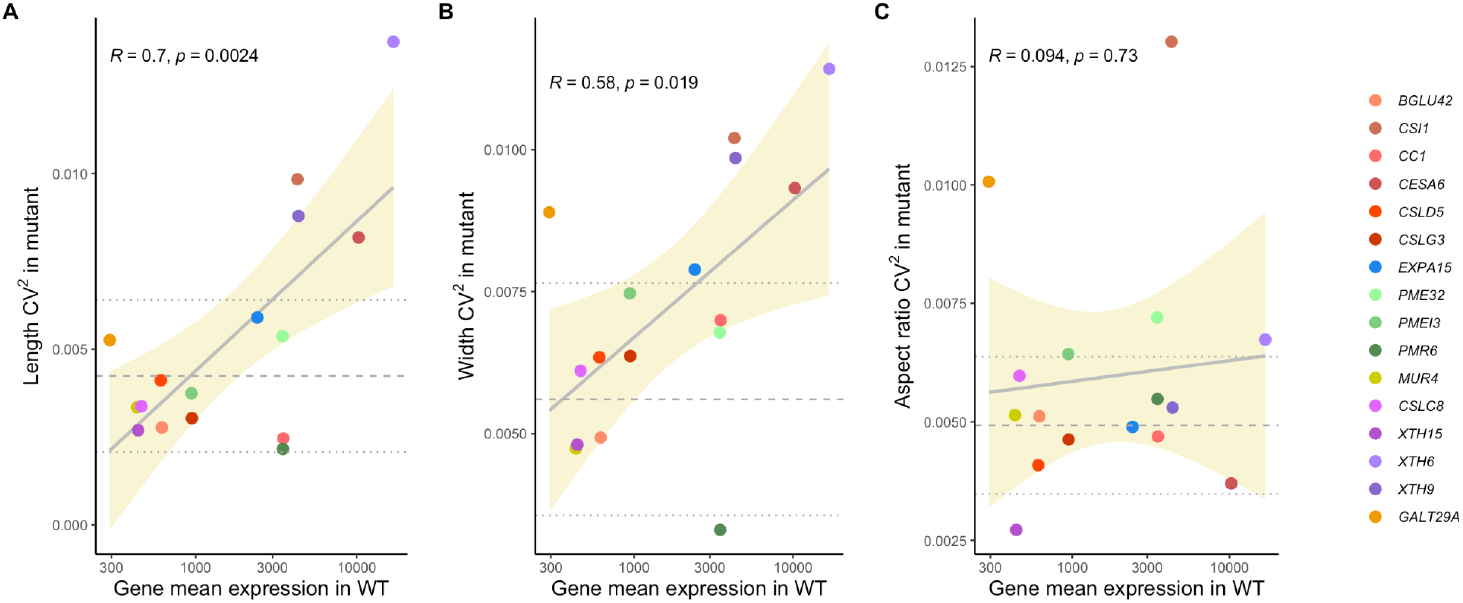
Squared coefficient of variation of sepal length (A), width (B) and aspect ratio (C) in knockout mutants against mean gene expression of the corresponding mutant genes in wild-type (WT) Col-0 plants. Each point corresponds to one gene. Pearson correlation coefficient R and p-values are shown on the top in each plot. Gray solid line shows linear model adjustment with standard error in tan shade. Gray dashed lines correspond to average CV^2^ in length, width and aspect ratio measurements in WT Col-0 control batches with dotted lines showing average plus/minus one standard deviation.

We performed additional tests to ensure the strength of our results. To confirm that sample size was not a confounding factor given its difference across our mutant dataset (from 39 to 90; Supplementary Table 3), we examined the correlation between parameter CV^2^ and sample size over all mutants and only found a marginally significant correlation value for Aspect ratio CV^2^. In addition, we performed 1000 replicates of subsampling experiments (see Materials and methods) obtaining R and p-values for 1000 correlation tests for each of our four parameters. Correlation p-values for Area CV^2^ and Length CV^2^ were below 0.05 (significant) in 100% of replicates and correlations for Width CV^2^ were significant in 47% of replicates. We then performed leave x-out experiments with x = {1, 2, 3} and calculated R and p-values for all possible combinations (see Materials and methods). Area CV^2^ and Length CV^2^ correlations with gene expression level are resilient to leaving 1, 2 and 3 mutants out, while many correlations for Width CV^2^ lose significance with the removal of 2 and 3 mutants (Figure 3). Through these experiments we observed that *PMR6* was not only an outlier, but that its presence affected the resilience of our main finding importantly. We hence proceeded to repeat the leave x-out experiments but after having completely removed *PMR6* from the data. These leave x-out experiments with x = {*pmr6*+1, *pmr6*+2, *pmr6*+3} reveal striking resilience of our results, with all correlations tested being below the 0.05 significance threshold and most of them depicting p-values below the one obtained for the complete set of mutants despite smaller sample sizes in the leave x-out experiments (Figure 3). These validations confirm that the expression level of our set of cell-wall related genes in wild type correlates with variability (and not average effect) in sepal area, length and width of the corresponding mutant.

**Figure 3.**
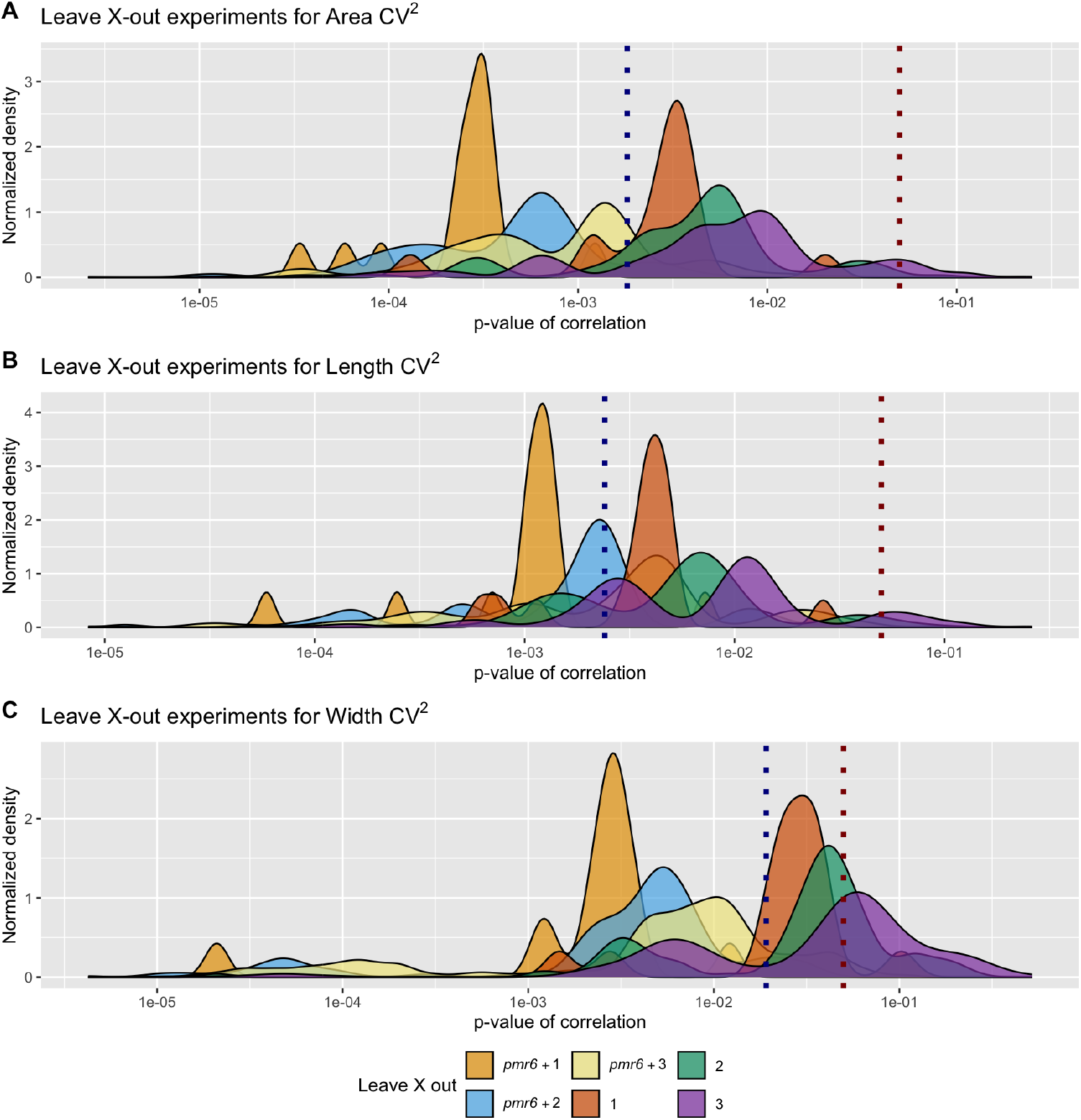
Leave x-out experiments for Area (A), Length (B) and Width (C) squared coefficient of variation (CV^2^) values. Each plot shows the distribution of p-values for all correlations tested within each of the six leave x-out experiments with x={1, 2, 3, *pmr6*+1, *pmr6*+2, *pmr6*+3}. The density curves are normalized to account for the different number of combinations tested in each experiment so that all areas under the curves are equal. Vertical dotted lines correspond to the p-value for the original correlation with the 16 mutants (dark blue) and to the significance threshold of p-value = 0.05 (dark red).

## Discussion

Although several signaling pathways regulating organ size and shape have been identified (reviewed in Powell & Lenhard, 2012), the mechanisms behind morphological robustness have remained elusive. More recently, a screen for mutants that disrupted the robustness of sepal size and shape led to the identification of genes involved in reproducibility of sepal morphogenesis (Hong et al., 2016; Zhu et al., 2020; Xu et al., 2024). This work revealed that spatio-temporal averaging of cellular variability, precise timing of organ initiation, and growth balance between cell layers, are required for precision in organ size and shape. The mechanisms underlying organ robustness, however, remain largely enigmatic.

Robustness of organ size and shape is thought to be the result of the complex interaction of genes within gene regulatory networks and environmental cues. Gene expression variability can be considered, *a priori*, as a factor affecting robustness. Hong et al. (2016) reported that cell-to-cell variability of growth is required for robustness of sepal shape and size, opening the question of the role of variability in gene expression in this process. Trinh et al. (2023) found that an increase in variability of gene expression impaired robustness of sepal shape and size. To further assess the link between gene expression variability and robustness, we have evaluated sepal size and shape in cell-wall related mutants. Therefore, we used mutants, not to search for candidate genes involved in size and shape determination, but to test the way in which knockout mutants of genes putatively involved in robustness, such as cell-wall related genes, affect sepal size and shape. Our results show that knockouts of highly expressed genes are associated with an increased phenotypic variability. In particular, highly expressed cell-wall related genes in wild-type plants exhibit larger morphological variation when knocked out than less expressed genes, although levels of active proteins do not necessarily correlate with RNA levels.

We can speculate that the effect caused by knocking out highly expressed cell-wall genes is more difficult to buffer, compared to lowly expressed cell-wall genes. There is knowledge about capacitors of phenotypic variation, such as heat-shock protein HSP90 (Queitsch et al., 2002), which limits the manifestation of cryptic genetic variation allowing for developmental stability (Sangster et al., 2008). Additionally, the effect of the mutation of a gene can be compensated by changes in the expression of other genes (e.g., Sénéchal et al., 2015; Hocq et al., 2020), possibly masking the mutation. Our work suggests that the function of genes can be revealed by analyzing variability of mutant phenotype instead of their average phenotype. Our observation that cell-wall related genes appear to be involved in buffering mechanisms that allow for developmental robustness, adds to evidence of cell-wall related genes being important for organ size and proportions (Weiss et al., 2005).

The set of mutants used in this study includes genes related to cellulose, hemicellulose, and pectin (Table 1). The selection of these genes was independent from the transcriptional study of Hartasánchez et al. (2023). Among the mutants analyzed, *mur4* has the largest effect in area, more than twice as that observed for *xth6* (Figure 1B & C). However, *xth6* shows five times more variability in sepal area than *mur4* (Figure 1D & E). Although each of the corresponding mutated genes is involved in a different metabolic process, namely, galactose for *MUR4* and xyloglucan for *XTH6*, finding a mechanistic explanation for this difference is not trivial. By comparing the *xth6* phenotype with other genes involved in xyloglucan metabolic processes, such as *XTH9* and *XTH15*, we observe that among these three genes, there is a negative correlation between Area CV^2^ and gene expression CV^2^ in wild type (Figure 1D, purple points) and a positive correlation between Area CV^2^ in the mutant and mean gene expression in wild type (Figure 1E, purple points). Whereas the former correlation is lost when including the other mutants, the latter holds and is statistically significant. This implies that regardless of the metabolic process any of these genes is involved in, the gene’s average expression in wild type seems to be an important parameter determining the variability in the effect of that gene’s knockdown mutation.

Testing for the resilience of our finding by performing leave x-out experiments revealed that the correlations for Area CV^2^ and Length CV^2^ were still significant after removal of one and two mutants, but less so for Width CV^2^. These tests also confirmed that *PMR6* was not only an outlier as could be noted by eye inspection of Figure 1 and Figure 2, but that its presence affected the resilience of the results importantly. Regarding the reasons why the *PMR6* gene could be an outlier, we might speculate that it is not directly involved in developmental robustness, as opposed to the other genes. *PMR6* is a pectate-lyase that makes Arabidopsis susceptible to powdery mildew (Vogel et al. 2002). It appears that *pmr6* resistance is not due to the activation of known host defense pathways but rather a novel form of resistance due to the loss of a gene required for a compatible interaction. Its exact function is not known, yet, it clearly affects cell wall composition and has pleiotropic effects on the plant (Vogel et al., 2002), so there is no clear reason why we would rule out the involvement of this gene in sepal development or robust development. However, by repeating the leave x-out experiments, but previously removing the *PMR6* gene from our list, the resilience of the correlation between variability in mutant size and gene expression in wild type is drastically increased (Figure 3).

Regarding the other genes found within our list, for example those involved in cellulose synthesis, we might also speculate about their role in developmental robustness. In mutants which have an affected cellulose synthesis, mechanical stresses might be created, setting off well known cell wall signaling events and potentially leading to transient growth cessation. Cytoskeletal interactions might also be involved in mechanical stress responses, involving proteins linking the cellulose synthase complex to cortical microtubules such as CSI1 (Mollier et al., 2023) and CC1 (Endler et al., 2015). However, the number of mutants investigated is too small to draw strong conclusions about the relation between gene function and phenotypic variability of the mutant. We do not know whether this phenotypic variability is a direct effect of cell wall modifications or involves feedback from the cell wall on cell growth, through cell wall integrity sensing and mechanosensing, for instance. Finally, we have only characterised the transcriptional regulation of robustness, discarding the potential role of signals (reactive oxygen species, calcium, pH, etc.) that directly affect cell wall properties.

Nevertheless, this work contributes to the understanding of how gene regulatory networks are related to developmental robustness. It opens questions regarding the role played by the level of expression of cell wall genes in the robustness of sepal size. Our results point towards genes with higher expression levels being more relevant, not for morphology, but for morphological robustness.

## Acknowledgements

This work was funded by the French National Research Agency (ANR) through a European ERA-NET Coordinating Action in Plant Sciences (ERA-CAPS) grant (Grant No. ANR-17-CAPS-0002-01 V-Morph) and through a direct grant (Grant No. ANR-17-CE20-0023-02 WALLMIME) by Fond de Recherche ENS de Lyon (Projet émergent FLORIVAR), by BAP INRAE (Projet FLORIVAR), and by a PhD fellowship from ENS de Lyon (to MD). We thank Annamaria Kiss and Marina Brasó-Vives for their help in early stages of this project. We thank Fabien Sénéchal and Jérôme Pelloux for providing the *pme32-1* seeds. Author contributions: Conceptualization - DAH, FM, AB; Methodology - DAH, MD, FM, AB; Software - DAH; Validation - DAH, MD, ND; Formal Analysis - DAH, MD; Investigation - DAH, MD, ND; Data Curation - DAH, MD, AB; Writing / Original Draft Preparation - DAH, FM, AB; Writing / Review & Editing - DAH, FM, AB; Visualization - DAH; Supervision - FM, AB; Project Administration - FM, AB; Funding Acquisition - FM, AB.

## Data availability

Raw data for mutant sepal measurements and their corresponding controls; Supplementary Tables 1-5; and the R scripts to analyze the raw data, to produce Supplementary Tables 2-5 and Figures 1 & 2 are publicly available at Zenodo (https://doi.org/10.5281/zenodo.13918461).

## References

Alonso, J. M., Stepanova, A. N., Leisse, T. J., Kim, C. J., Chen, H., Shinn, P., Stevenson, D. K., Zimmerman, J., Barajas, P., Cheuk, R., Gadrinab, C., Heller, C., Jeske, A., Koesema, E., Meyers, C. C., Parker, H., Prednis, L., Ansari, Y., Choy, N., Deen, H., … Ecker, J. R. (2003). Genome-wide insertional mutagenesis of Arabidopsis thaliana. Science, 301(5633), 653–657. 10.1126/science.1086391

Alvarez-Buylla, E. R., Chaos, A., Aldana, M., Benítez, M., Cortes-Poza, Y., Espinosa-Soto, C., Hartasánchez, D. A., Lotto, R. B., Malkin, D., Escalera Santos, G. J., & Padilla-Longoria, P. (2008). Floral morphogenesis: stochastic explorations of a gene network epigenetic landscape. PloS One, 3(11), e3626. 10.1371/journal.pone.0003626

Araújo, I. S., Pietsch, J. M., Keizer, E. M., Greese, B., Balkunde, R., Fleck, C., & Hülskamp, M. (2017). Stochastic gene expression in Arabidopsis thaliana. Nature Communications, 8(1), 2132. 10.1038/s41467-017-02285-7

Bringmann, M., Li, E., Sampathkumar, A., Kocabek, T., Hauser, M. T., & Persson, S. (2012). POM-POM2/cellulose synthase interacting1 is essential for the functional association of cellulose synthase and microtubules in Arabidopsis. The Plant Cell, 24(1), 163–177. 10.1105/tpc.111.093575

Cortijo, S., Aydin, Z., Ahnert, S., & Locke, J. C. (2019). Widespread inter-individual gene expression variability in Arabidopsis thaliana. Molecular Systems Biology, 15(1), e8591. 10.15252/msb.20188591

Desnos, T., Orbović, V., Bellini, C., Kronenberger, J., Caboche, M., Traas, J., & Höfte, H. (1996). Procuste1 mutants identify two distinct genetic pathways controlling hypocotyl cell elongation, respectively in dark- and light-grown Arabidopsis seedlings. Development, 122(2), 683–693. 10.1242/dev.122.2.683

Dumond, M. (2017). From cellular variability to shape reproducibility: mechanics and morphogenesis of Arabidopsis thaliana sepal. Université de Lyon, Lyon. https://tel.archives-ouvertes.fr/tel-01650126

Elowitz, M. B., Levine, A. J., Siggia, E. D., & Swain, P. S. (2002). Stochastic gene expression in a single cell. Science, 297(5584), 1183–1186. 10.1126/science.1070919

Endler, A., Kesten, C., Schneider, R., Zhang, Y., Ivakov, A., Froehlich, A., Funke, N., & Persson, S. (2015). A Mechanism for Sustained Cellulose Synthesis during Salt Stress. Cell, 162(6), 1353–1364. 10.1016/j.cell.2015.08.028

Félix, M. A., & Barkoulas, M. (2015). Pervasive robustness in biological systems. Nature Reviews Genetics, 16(8), 483–496. 10.1038/nrg3949

Hartasánchez, D. A., Kiss, A., Battu, V., Soraru, C., Delgado-Vaquera, A., Massinon, F., Brasó-Vives, M., Mollier, C., Martin-Magniette, M., Boudaoud, A., & Monéger, F. (2023). Expression of cell-wall related genes is highly variable and correlates with sepal morphology. Peer Community Journal, 3: e93. 10.24072/pcjournal.327

Hocq, L., Guinand, S., Habrylo, O., Voxeur, A., Tabi, W., Safran, J., Fournet, F., Domon, J. M., Mollet, J. C., Pilard, S., Pau-Roblot, C., Lehner, A., Pelloux, J., & Lefebvre, V. (2020). The exogenous application of AtPGLR, an endo-polygalacturonase, triggers pollen tube burst and repair. The Plant Journal, 103(2), 617–633. 10.1111/tpj.14753

Hong, L., Dumond, M., Tsugawa, S., Sapala, A., Routier-Kierzkowska, A. L., Zhou, Y., Chen, C., Kiss, A., Zhu, M., Hamant, O., Smith, R. S., Komatsuzaki, T., Li, C. B., Boudaoud, A., & Roeder, A. H. (2016). Variable Cell Growth Yields Reproducible OrganDevelopment through Spatiotemporal Averaging. Developmental Cell, 38(1), 15–32. 10.1016/j.devcel.2016.06.016

Joseph, B., Corwin, J. A., & Kliebenstein, D. J. (2015). Genetic variation in the nuclear and organellar genomes modulates stochastic variation in the metabolome, growth, and defense. PLoS Genetics, 11(1), e1004779. 10.1371/journal.pgen.1004779

Lachowiec, J., Queitsch, C., & Kliebenstein, D. J. (2016). Molecular mechanisms governing differential robustness of development and environmental responses in plants. Annals of Botany, 117(5), 795–809. 10.1093/aob/mcv151

Liu, J., Frochaux, M., Gardeux, V., Deplancke, B., & Robinson-Rechavi, M. (2020). Inter-embryo gene expression variability recapitulates the hourglass pattern of evo-devo. BMC Biology, 18(1), 129. 10.1186/s12915-020-00842-z

McElver, J., Tzafrir, I., Aux, G., Rogers, R., Ashby, C., Smith, K., Thomas, C., Schetter, A., Zhou, Q., Cushman, M. A., Tossberg, J., Nickle, T., Levin, J. Z., Law, M., Meinke, D., & Patton, D. (2001). Insertional mutagenesis of genes required for seed development in Arabidopsis thaliana. Genetics, 159(4), 1751–1763. 10.1093/genetics/159.4.1751

Meyer, H. M., Teles, J., Formosa-Jordan, P., Refahi, Y., San-Bento, R., Ingram, G., Jönsson, H., Locke, J. C., & Roeder, A. H. (2017). Fluctuations of the transcription factor ATML1 generate the pattern of giant cells in the Arabidopsis sepal. eLife, 6, e19131. 10.7554/eLife.19131

Mollier, C., Skrzydeł, J., Borowska-Wykręt, D., Majda, M., Bayle, V., Battu, V., Totozafy, J. C., Dulski, M., Fruleux, A., Wrzalik, R., Mouille, G., Smith, R. S., Monéger, F., Kwiatkowska, D., & Boudaoud, A. (2023). Spatial consistency of cell growth direction during organ morphogenesis requires CELLULOSE SYNTHASE INTERACTIVE1. Cell reports, 42(7), 112689. 10.1016/j.celrep.2023.112689

O’Malley R.C., Barragan C.C., Ecker J.R. (2015) A User’s Guide to the Arabidopsis T-DNA Insertion Mutant Collections. In: Alonso J., Stepanova A. (eds) Plant Functional Genomics. Methods in Molecular Biology, 1284. Humana Press, New York, NY. 10.1007/978-1-4939-2444-8_16

Powell, A. E., & Lenhard, M. (2012). Control of organ size in plants. Current Biology, 22(9), R360–R367. 10.1016/j.cub.2012.02.010

Queitsch, C., Sangster, T. A., & Lindquist, S. (2002). Hsp90 as a capacitor of phenotypic variation. Nature, 417(6889), 618–624. 10.1038/nature749

R Core Team (2020). R: A language and environment for statistical computing. R Foundation for Statistical Computing, Vienna, Austria. URL https://www.R-project.org/

Sangster, T. A., Salathia, N., Lee, H. N., Watanabe, E., Schellenberg, K., Morneau, K., Wang, H., Undurraga, S., Queitsch, C., & Lindquist, S. (2008). HSP90-buffered genetic variation is common in Arabidopsis thaliana. Proceedings of the National Academy of Sciences of the United States of America, 105(8), 2969–2974. 10.1073/pnas.0712210105

Schmalhausen, I. I. (1949). Factors of evolution: the theory of stabilizing selection. Univ. of Chicago Press, Chicago, IL.

Schmid, M., Davison, T.S., Henz, S.R., Pape, U.J., Demar, M., Vingron, M., Schölkopf, B., Weigel, D., and Lohmann, J.U. (2005). A gene expression map of Arabidopsis thaliana development. Nature Genetics, 37, 501–506. 10.1038/ng1543.

Sénéchal, F. (2013). Rôles des pectines méthylestérases (PMEs) dans le développement chez Arabidopsis thaliana. Étude de leur régulation par les inhibiteurs (PMEIs) et protéases de type subtilisines (SBTs). Université de Picardie Jules Verne, Amiens.

Sénéchal, F., Mareck, A., Marcelo, P., Lerouge, P., & Pelloux, J. (2015). Arabidopsis PME17 Activity can be Controlled by Pectin Methylesterase Inhibitor4. Plant Signaling & Behavior, 10(2), e983351. 10.4161/15592324.2014.983351

Smyth, D. R., Bowman, J. L., & Meyerowitz, E. M. (1990). Early flower development in Arabidopsis. The Plant Cell, 2(8), 755–767. 10.1105/tpc.2.8.755

Trinh, D.-C., Martin, M., Bald, L., Maizel, A., Trehin, C., and Hamant, O. (2023). Increased gene expression variability hinders the formation of regional mechanical conflicts leading to reduced organ shape robustness. Proceedings of the National Academy of Sciences of the United States of America, 120, e2302441120. 10.1073/pnas.2302441120

Van Rossum, G., & Drake, F. L. (2009). Python 3 Reference Manual. Scotts Valley, CA: CreateSpace. https://docs.python.org/3/reference/index.html

Vogel, J. P., Raab, T. K., Schiff, C., & Somerville, S. C. (2002). PMR6, a pectate lyase-like gene required for powdery mildew susceptibility in Arabidopsis. The Plant cell, 14(9), 2095–2106. 10.1105/tpc.003509

Waddington, C. H. (1953). Assimilation of an acquired character. Evolution, 7(2): 118–126. 10.2307/2405747

Waddington, C. H. (1959). Canalization of development and genetic assimilation of acquired characters. Nature, 183(4676), 1654–1655. 10.1038/1831654a0

Weiss, J., Delgado-Benarroch, L., & Egea-Cortines, M. (2005). Genetic control of floral size and proportions. The International Journal of Developmental Biology, 49(5-6), 513–525. 10.1387/ijdb.051998jw

Xu, S., He, X., Trinh, D.-C., Zhang, X., Wu, X., Qiu, D., Zhou, M., Xiang, D., Roeder, A.H.K., Hamant, O., and Hong, L. (2024). A 3-component module maintains sepal flatness in Arabidopsis. Current Biology, 34, 4007-4020.e4. 10.1016/j.cub.2024.07.066

Zalts, H., & Yanai, I. (2017). Developmental constraints shape the evolution of the nematode mid-developmental transition. Nature Ecology & Evolution, 1(5), 113. 10.1038/s41559-017-0113

Zhu, M., Chen, W., Mirabet, V., Hong, L., Bovio, S., Strauss, S., Schwarz, E. M., Tsugawa, S., Wang, Z., Smith, R. S., Li, C. B., Hamant, O., Boudaoud, A., & Roeder, A. (2020). Robust organ size requires robust timing of initiation orchestrated by focused auxin and cytokinin signalling. Nature Plants, 6(6), 686–698. 10.1038/s41477-020-0666-7

